# Analysis of deep learning methods for blind protein contact prediction in CASP12

**DOI:** 10.1101/181586

**Authors:** Sheng Wang, Siqi Sun, Jinbo Xu

## Abstract

Here we present the results of protein contact prediction achieved in CASP12 by our RaptorX-Contact server, which is an early implementation of our deep learning method for contact prediction. On a set of 38 free-modeling target domains with a median family size of around 58 effective sequences, our server obtained an average top L/5 long- and medium-range contact accuracy of 47% and 44%, respectively (L=length). A more advanced implementation has an average accuracy of 59% and 57%, respectively. Our deep learning method formulates contact prediction as an image pixel-level labeling problem and simultaneously predicts all residue pairs of a protein using a combination of two deep residual neural networks, taking as input the residue conservation information, predicted secondary structure and solvent accessibility, contact potential, and co-evolution information. Our approach differs from existing methods mainly in (1) formulating contact prediction as a pixel-level image labeling problem instead of an image-level classification problem; (2) simultaneously predicting all contacts of an individual protein to make effective use of contact occurrence patterns; and (3) integrating both 1D and 2D deep convolutional neural networks to effectively learn complex sequence-structure relationship including high-order residue correlation. This paper discusses the RaptorX-Contact pipeline, both contact prediction and contact-based folding results, and finally the strength and weakness of our method.

## Introduction

Ab initio protein folding or de novo structure prediction is one of most challenging problems in computational biology. The popular fragment assembly method mainly works well on some small proteins. Recent progress has indicated that long-range contacts are very helpful for structure prediction [1] and that direct co-evolution analysis may correctly predict some long-range contacts for proteins with a very large number of sequence homologs [2, 3]. However, soluble proteins with that many sequence homologs likely have similar solved structures in PDB and thus, may be modelled by template-based methods. For proteins with few sequence homologs, pure co-evolution methods such as CCMpred[4], PSICOV[5], Evfold[6], Gremlin[7], and CoinDCA[8] do not fare well and their predictions are not very helpful to ab initio folding. Supervised learning such as PhyCMAP[9], DNCON[10] and SVMSEQ[11] predicts contacts using a variety of protein features, on average outperforming pure co-evolution methods on proteins with few sequence homologs.

In CASP11, MetaPSICOV[12] stands out as an excellent contact predictor. It predicts contacts by combining direct co-evolution information generated by several statistical methods and a few “classic” protein features such as sequence profile and predicted local structure properties. MetaPSICOV embraces the advantage of both co-evolution analysis and supervised learning, but its accuracy on proteins without many sequence homologs is still unsatisfactory. To further improve contact prediction especially for small-sized protein families, we have developed a deep learning method [13] that can effectively integrate both “classic” protein features and co-evolution information. Our deep learning method is good at learning contact occurrence patterns and predicting protein-like contact maps, and thus significantly improves contact prediction accuracy. Contact occurrence patterns describe multi-body residue correlation learned from native structures, which is orthogonal to pairwise co-evolution information (extracted from sequences) and the “classic” protein features (e.g., sequence profile and predicted local structure properties).

In this paper, we analyze the deep learning method we developed during CASP12. Since we were developing our method during the whole CASP12 season, the deep models we used in CASP12 varied as we gradually improved them through the end of CASP12. We started by formulating contact prediction as an image-level classification problem and then switched the formulation to a pixel-level labeling problem. Pixel-level image labeling refers to each pixel in an image having a label to be predicted whereas image-level classification means that the whole image has only one label to be predicted (e.g., face recognition). We also gradually added more convolution and batch normalization[14] layers to our deep models and optimized our training algorithm to yield better model parameter estimation. Minor improvement includes searching for sequence homologs using different E-values and splitting a multi-domain protein into domains. Here we focus on the major ideas we used in CASP12, discuss our performance and analyze the strengths and weaknesses of our approach. We also examine several interesting free modeling cases and discuss the models we have built and can build using our predicted contacts. Finally, we highlight our views on the future development of contact prediction and its challenge.

## Materials and Methods

Following the CASP definition, we say two residues form a contact if in the native structure, the distance of their C_β_ atoms is less than 8Å. Contacts in a protein are not randomly distributed, instead they are sparse and form some patterns.

### Multiple sequence alignment (MSA) and input features

We generate MSAs in three ways. To save time in generating features for the training proteins, we used the MSAs already in our template-based RaptorX server[15]. These MSAs were generated before 2016 by the buildAli.pl program in the HHpred package[16], which calls PSI-BLAST (5 iterations and E-value=0.001) to find sequence homologs from an old NR database and then build MSAs. For CASP12 targets, initially we generated one MSA for each target by running HHblits[17] with 3 iterations and E-value set to 0.001 on the unitprot20 library released in February 2016. At the late stage of CASP12, for each test target we generated four different MSAs by running HHblits with 3 iterations and E-value set to 0.001 and 1, respectively, to search through the uniprot20 library released in November 2015 and February 2016, respectively. That is, for each target we generated 4 sets of input features and accordingly 4 different contact predictions, which are then averaged to obtain the final prediction. Our in-house test shows that using 4 rather than 1 MSAs may yield 1-2% accuracy improvement in contact prediction.

From each individual MSA, we derive two types of protein features: sequential features and pairwise features. Sequential features include sequence profile and secondary structure and solvent accessibility predicted by our RaptorX-Property[18]. Pairwise features include mutual information, pairwise contact potential and direct co-evolution strength calculated by CCMpred. Whereas MetaPSICOV uses three co-evolution analysis tools to generate direct co-evolution information, we employed only CCMpred, which runs very fast on GPUs yet has very good accuracy.

### Pixel-level labeling formulation vs. image-level classification formulation

Pixel-level image labeling refers to each pixel in an image having a label whereas in image-level classification, the whole image has only one label (e.g., face recognition). By formulating contact prediction as a pixel-level image labeling problem, we simultaneously predict the labels of all the entries in a contact matrix without splitting this contact matrix into small submatrices. By doing so, the prediction error at one residue pair may be fed back to all the other residue pairs through back propagation (in training). Such a formulation also makes it easy to exploit high-order residue correlation or contact correlation. By contrast, existing methods such as MetaPSICOV formulate contact prediction as an image-level classification problem by separating the prediction of one residue pair from the others. For each residue pair under prediction, these methods extract a submatrix centering around this residue pair and predict one label for this submatrix (e.g., the center residue pair being in contact or not in contact). Since the submatrix is cut off from the original contact matrix, the prediction error at its center residue pair cannot be effectively fed back to the other residue pairs. Such a formulation also makes it challenging to directly model contact correlation. To predict the complete contact matrix of a protein with L residues, these other methods need to predict the labels of L*(L-6)/2 submatrices one-by-one while our method directly works on the whole contact matrix. This implies that pixel-level formulation is also computationally more efficient than image-level formulation (given the same network architecture and the same number of residue pairs to be predicted).

### Deep learning models

We treat a protein contact map as an image of L×L where L is sequence length and formulate contact prediction as a pixel-level image labeling problem. This is different from the formulation employed by MetaPSICOV, which formulates contact prediction as an image-level classification problem. However, we cannot directly apply the deep models developed for image labeling to contact prediction since proteins have more complex features and the ratio of contacts (i.e., positive labels) is very small (<2%). Instead we developed a deep learning model by concatenating two deep residual neural networks.

As shown in Fig. 1(A), the first residual network conducts a 1-dimensional (1D) convolutional transformation of sequential features to capture long-range sequential context of a residue. Its output is converted to a 2-dimensional (2D) matrix by an operation similar to outer product and then fed into the 2^nd^ residual network together with the original pairwise features. The 2^nd^ residual network conducts 2D convolutional transformation of its input to capture long-range 2D context of a residue pair. Finally, the output of the 2^nd^ network is fed into logistic regression, which predicts the probability of any two residues in a contact.

**Figure 1.**
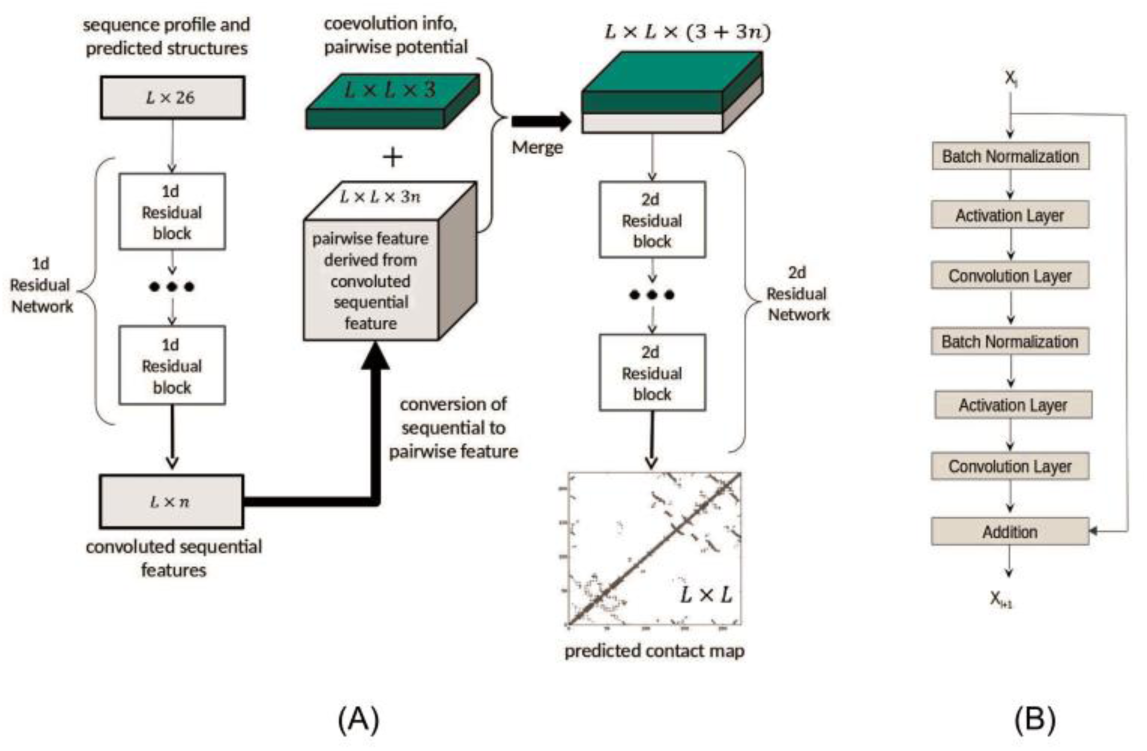
(A) The overall network architecture of the deep learning model. Meanwhile, L is protein sequence length and n is the number of hidden neurons in the last 1D convolutional layer. (B) The internal structure of a residual block with X_l_ and X_l+1_ being input and output, respectively.

Each residual network is composed of some residual blocks, each block in turn consisting of 2 batch normalization layers, 2 convolution layers and 2 ReLU activation layers (Fig. 1(B)). In the first residual network, X_l_ and X_l+1_ represent sequential features and have dimension L×n _l_ and L×n _l+1_, respectively, where n_l_ (n_l+1_) is the number of hidden neurons at each residue. In the 2nd residual network, X_l_ and X_l+1_ represent pairwise features and have dimension L×L×n _l_ and L×L×n _l+1_, respectively, where n_l_ (n_l+1_) is the number of hidden neurons at one residue pair. The filter size (i.e., window size) used by a 1D convolution layer is 15 or 17 while that used by a 2D convolution layer is 3×3 or 5×5. We fix the depth (i.e., the number of convolution layers) of the first residual network to 6 and vary the depth of the second network. Each 1D convolutional layer has around 50 hidden neurons. With ∼50 hidden neurons at each of the 1D convolutional layers, 55-75 hidden neurons at of the 2D convolutional layers, and 50-60 convolution layers for the 2^nd^ network, our model can yield pretty good performance.

Our deep learning method is unique in the following aspects. First, the formulation of pixel-level labeling is different from the image-level classification formulation employed by many existing methods such as MetaPSICOV. Second, our model employs a combination of two deep residual neural networks, which has not been applied to contact prediction before. Third, we predict all contacts of a protein simultaneously, as opposed to existing supervised methods that predict contacts one by one. By simultaneous prediction, we can easily learn contact occurrence patterns from protein structures so that our predicted contact maps are more protein-like. By contrast, current co-evolution methods focus only on pairwise relationship extracted from sequences. Finally, our deep model learn knowledge from thousands of non-redundant protein families, as opposed to current co-evolution methods that use information only in a single protein family.

### Model training and validation

We train our models by maximum-likelihood with L_2_-norm regularization. The objective function to be minimized in training our deep model is the negative log-likelihood averaged per residue pair plus the L2 norm of model parameters multiplied by the regularization factor. As long as its value is between 0.005 and 0.0001, the regularization factor does not impact prediction accuracy much. We train our deep models using minibatches and run our training algorithm for only 20 epochs. Each epoch scans through all the training proteins once and each minibatch has several proteins. Proteins in different minibatches may have different lengths while proteins in the same minibatch are forced to have the same length by zero padding. A stochastic gradient descent algorithm is used to minimize the objective function. The whole algorithm is implemented with Theano and runs on a GPU card. It took about one week to train a single deep model with 6000-7000 training proteins.

#### Training and test data

We have self-tested our method using the 150 Pfam families, the 105 CASP11 test proteins, 398 non-redundant membrane proteins and 76 hard CAMEO test proteins released from 10/17/2015 to 04/09/2016 (see [13] for the detailed results of these test sets). Our training set is a subset of PDB25 created in 2015, in which no two proteins share more than 25% sequence identity. To remove redundancy, we also exclude the proteins having a BLAST E-value <0.1 with any of the test proteins. In total there are ∼6300 training proteins plus 400 validation proteins, from which we have trained several models, which are then averaged to produce the final model.

#### Calculating the number of effective sequence homologs

Meff measures the amount of homologous information in an MSA. It can be interpreted as the number of non-redundant (or effective) sequence homologs in an MSA when 70% sequence identity is used as cutoff. To calculate Meff, we first calculate the sequence identity between any two proteins in the MSA. Let a binary variable S_ij_ denote the similarity between two protein sequences i and j. S_ij_ is equal to 1 if and only if the sequence identity between i and j is at least 70%. For a protein i, we calculate the sum of S_ij_ over all the proteins (including itself) in the MSA and denote it as S_i_. Finally, we calculate Meff as the sum of 1/S_i_ over all the protein sequences in this MSA.

## Results

### Contact prediction accuracy in CASP12

Table 1 summarizes the average contact prediction accuracy of our method RaptorX-submit and a few others on the 38 CASP12 FM targets. See Supplementary File 1 for the accuracy on each target. The top L/k (k=1, 2, 5, 10) contact prediction accuracy is the number of correctly-predicted contacts divided by L/k independent of how many native contacts exist. In addition to the predicted contacts submitted to CASP12, we also evaluate our deep learning method (denoted as RaptorX-postdict) trained right after the CASP12 season (i.e., between August and September 2016). This deep model still uses the uniprot20 sequence database dated in February 2016 to generate MSAs. As mentioned before, we were gradually improving our deep models until the end of the CASP12 season. Therefore, by comparing our CASP12 submissions with the predictions from our model trained right after CASP12, we can gauge how much improvement can be achieved by a well-developed deep learning method. This comparison is fair since the underlying sequence database for MSA generation is unchanged and all the training proteins were publicly released in PDB before CASP12 started. As shown in Table 1, RaptorX-postdict is much better than RaptorX-submit, but the advantage is much smaller at the end of CASP12 (see Supplementary Files 1 and 2 for details). While RaptorX-postdict outperforms RaptorX-submit on most targets, RaptorX-postdict underperforms on two domains T0918-D1 and T0918-D3. This worse performance is due to that RaptorX-postdict employed 4 different MSAs to produce an average prediction. If only the MSA generated from uniprot20-2016 with E-value=0.001 is used, RaptorX-postdict has the same accuracy as RaptorX-submit on T0918-D1 and slightly better accuracy on T0918-D3. RaptorX-submit used only the MSA generated from uniprot20-2016 with E-value=0.001 on this target.

**Table 1.**
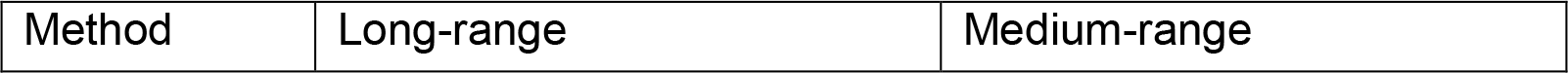
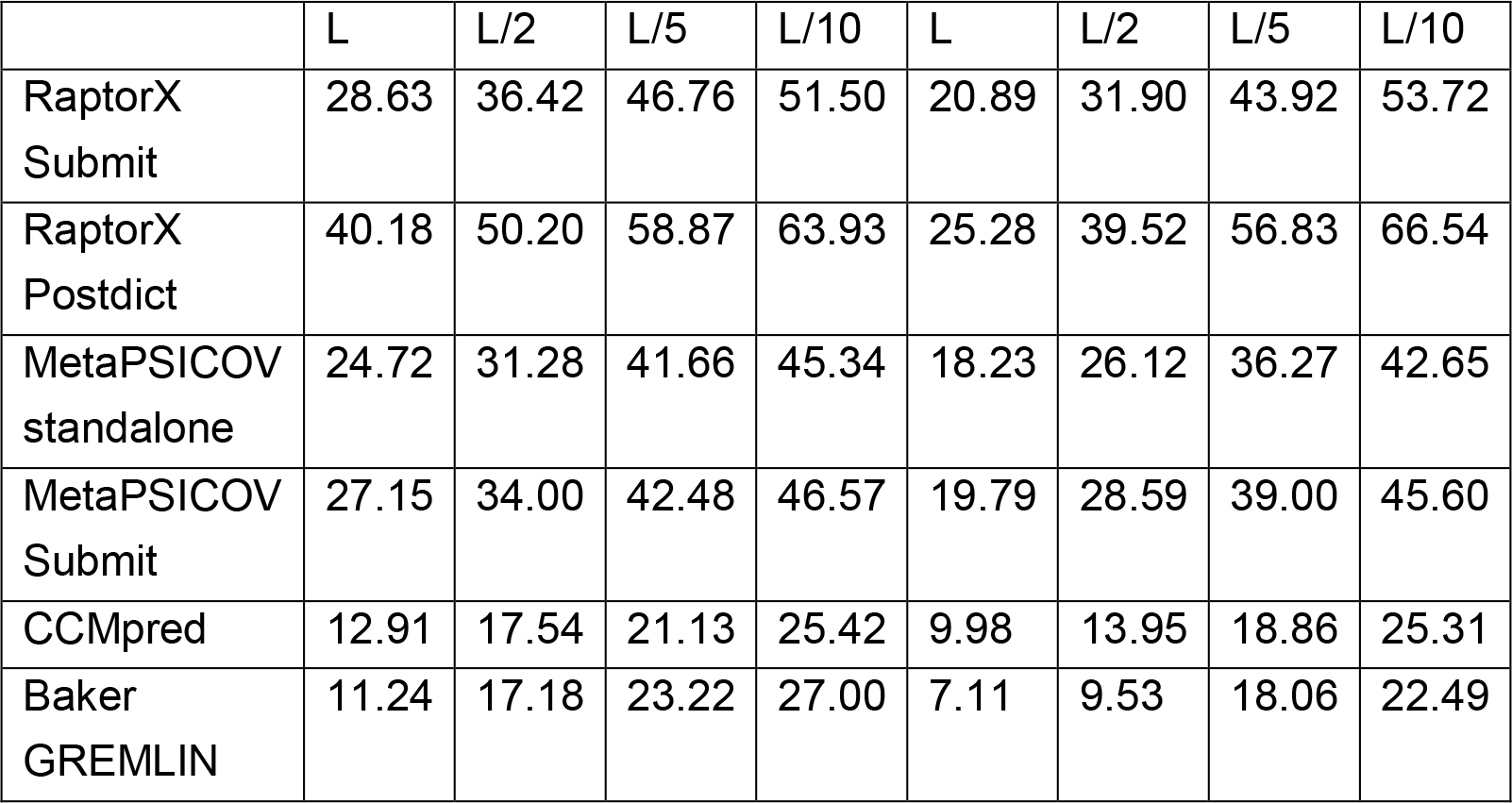
The average contact prediction accuracy of several methods on the CASP12 hard targets. See the main text for explanation and Supplementary File 1 for details.

In addition to the MetaPSICOV group in CASP12 (denoted as MetaPSICOV-submit), we also studied the performance of the standalone MetaPSICOV program publicly released by Jones group after CASP12 (denoted MetaPSICOV-standalone). The results of MetaPSICOV-submit were downloaded from the CASP12 web site and we ran MetaPSICOV-standalone locally using the same uniprot20 sequence database as our deep learning method. MetaPSICOV-submit has slightly better accuracy than MetaPSICOV-standalone. This may be because that our method of generating MSAs is not as good as that of the Jones group. We also examined the performance of CCMpred and Baker-GREMLIN, the state-of-the art pure co-evolution methods for contact prediction. We ran CCMpred locally with the same uniprot20 sequence database. The results of Baker-GREMLIN are downloaded from the CASP12 web site. CCMpred has similar long-range accuracy as GREMLIN, but much better medium-range accuracy.

In summary, our submitted predictions are slightly better than MetaPSICOV submissions, but our fully-implemented deep model trained right after CASP12 performs much better than MetaPSICOV, which is consistent with our previous results [13]. This indicates that our deep neural network indeed performs much better than the shallow neural network used by MetaPSICOV. The CASP12 results show that pure co-evolution methods do not work well on the CASP12 hard targets mainly because most of them do not have many sequence homologs. However, even for targets with more than 1000 effective sequence homologs such as T0866-D1, T0899-D1, T0905-D1, T0918-D1 and T0918-D3, MetaPSICOV and our deep learning method still greatly outperform CCMpred and GREMLIN (Table 2). This may confirm that MetaPSICOV and our method indeed introduce extra information (orthogonal to co-evolution information) for contact prediction.

**Table 2.** The top L/2 long-range contact prediction accuracy on five CASP12 hard targets with more than 1000 effective sequence homologs. Std represents standard deviation.

| Target | Length | Meff | CCMpred | Baker<br>GREMLIN | MetaPSICOV<br>submit | RaptorX<br>submit | RaptorX<br>postdict |
| --- | --- | --- | --- | --- | --- | --- | --- |
| T0866-D1 | 104 | 3383 | 88.46 | 76.92 | 96.15 | 84.62 | 94.23 |
| T0899-D1 | 259 | 1047 | 31.01 | 0.78 | 57.36 | 57.36 | 71.32 |
| T0905-D1 | 242 | 4967 | 52.07 | 31.40 | 77.69 | 76.86 | 82.64 |
| T0918-D1 | 108 | 2827 | 44.44 | 50.00 | 62.96 | 72.22 | 70.37 |
| T0918-D3 | 118 | 2853 | 37.29 | 45.76 | 57.63 | 74.58 | 64.41 |
| Average | 166 | 3015 | 50.65 | 40.97 | 70.36 | 73.13 | 76.59 |
| Std | 69.2 | 1255 | 20.17 | 24.91 | 14.86 | 8.92 | 10.61 |

### Contact-assisted folding accuracy

We examined the accuracy of 3D models built from our predicted contacts. During CASP12, we built 3D models by feeding our predicted secondary structure and top L predicted contacts to the CNS package [19], which builds 3D models by treating contacts as distance restraints and converting predicted secondary structure to backbone torsion angles. At the very late stage of CASP12, we revised our 3D modeling strategy. In particular, we first normalized the predicted contact probability values by their mean and standard deviation calculated on the whole contact matrix. Then we selected those top predicted contacts with normalized value ≥3 and fed them to CNS. We also added or removed some contacts to make the number of selected contacts between L and 3L. To evaluate this new protocol, we rebuilt 3D models from our submitted contact predictions for all the CASP12 targets to see how much we may improve 3D model quality. We denote such a method as RaptorX-rebuild. In addition, we also built 3D models for all CASP12 targets using the contacts predicted by RaptorX-postdict (i.e., the deep model trained right after CASP12). As a comparison, we examined the quality of the 3D models built from CCMpred- and MetaPSICOV-predicted contacts using the same contact selection and 3D model building protocols. We also evaluated the 3D models submitted to CASP12 by RaptorX-TBM (i.e., our template-based modeling server), Baker-server, Zhang-server, Baker-human and Zhang-human. The latter four used a much more sophisticated protocol (i.e., hybrid of template-based and template-free methods) to predict 3D models than us. It is also possible that the two human groups used information from other CASP12-participating servers.

As shown in Table 3, the 3D models submitted to CASP12 by RaptorX-submit (i.e., RaptorX-Contact group) are not very good although on average they are better than CCMpred- and slightly better than MetaPSICOV-derived 3D models. RaptorX-rebuild is slightly better than RaptorX-submit. That is, by using more than top L contacts submitted to CASP12, we can slightly improve 3D modeling accuracy. The underlying reason why RaptorX-rebuild is not significantly better than RaptorX-submit is because the accuracy of our submitted contacts is not good enough. This finding is consistent with our observation that by using the same contact selection and model building protocols to build 3D models from CCMpred-or MetaPSICOV-predicted contacts, we cannot improve 3D modeling accuracy much either over using only top L contacts. However, by using the contacts predicted from RaptorX-postdict, we can generate much better 3D models with quality better than RaptorX-TBM and comparable to the 3D models submitted by Baker and Zhang servers, although RaptorX-postdict makes use of only predicted contacts and secondary structure. Nevertheless, RaptorX-postdict is slightly worse than Baker and Zhang human groups. See Supplementary File 3 for the 3D modeling accuracy of each CASP12 hard target.

**Table 3.** The average quality (TMscore) of the 3D models built by different methods (groups) for the CASP12 hard targets. Top k (k=1, 5) means that for each target the best of top k 3D models is considered. See Supplementary File 3 for details.

| Group | Top 1 | Top 5 |
| --- | --- | --- |
| RaptorX-submit | 0.274 | 0.307 |
| RaptorX-rebuild | 0.285 | 0.322 |
| RaptorX-postdict | 0.354 | 0.397 |
| RaptorX-TBM | 0.320 | N/A |
| Baker-server | 0.326 | 0.370 |
| Zhang-server | 0.347 | 0.404 |
| Baker-human | 0.392 | 0.422 |
| Zhang-human | 0.375 | 0.420 |
| CCMpred | 0.216 | 0.235 |
| MetaPSICOV | 0.262 | 0.289 |

Our deep learning method can generate good models for several targets with very few sequence homologs. For example, RaptorX-postdict can generate 3D models with TMscore>0.6 for T0864-D1, T0869-D1 and T0904-D1. They have only 209, 19, and 34 effective sequence homologs, respectively. The 3D models generated by RaptorX-postdict for them have much better quality than RaptorX-TBM, Baker- and Zhang-server and Baker- and Zhang-human groups. In contrast, Baker-human generated 3D models with TMscore>0.6 for 4 targets, among which two have >800 effective sequence homologs and the other two have >3000 effective sequence homologs. This may imply that Baker-human did very well as long as pure co-evolution methods can provide some correctly-predicted contacts. RaptorX-postdict also generates reasonable 3D models (TMscore ∼0.5) for T0862-D1, T0863-D1, T0866-D1, T0870-D1, T0886-D1, T0886-D2, T0899-D1, T0905-D1, and T0915-D1. Four of these targets have fewer than 100 effective sequence homologs.

## Case Study

In this section, we study in detail three specific targets T0864-D1, T0869-D1 and T0904-D1 on which RaptorX-postdict performs well.

### T0864-D1

Table 4 shows that for this target our method produced much better contact prediction than CCMpred and MetaPSICOV, especially when top L contacts are evaluated. Specifically, the contact map predicted by our method has L long-range accuracy 69.1%, while that by CCMpred and MetaPSICOV has corresponding accuracy 20.3% and 36.2%, respectively. Fig. 2 visualizes the top L/2 predicted contacts of the three methods as compared to the native contact map. The best of top 5 3D model submitted by our contact server has TMscore 0.63 and RMSD 4.48Å. The best of top 5 models built by CNS from CCMpred- and MetaPSICOV-predicted contacts have TMscore 0.274 and 0.353, respectively. Fig. 3 shows the superimposition between the native structure and the 3D models generated by our method, CCMpred and MetaPSICOV, respectively. To examine the superimposition of our model with its native structure from various angles, please see http://raptorx.uchicago.edu/DeepAlign/63883537/.

**Table 4.** The long- and medium-range contact prediction accuracy of our method, CCMpred, MetaPSICOV (submit) and on the CASP12 target T0864-D1.

|  | Long-range accuracy |  |  |  | Medium-range accuracy |  |  |  |
| --- | --- | --- | --- | --- | --- | --- | --- | --- |
|  | L | L/2 | L/5 | L/10 | L | L/2 | L/5 | L/10 |
| Our method | 0.691 | 0.854 | 1.000 | 1.000 | 0.447 | 0.756 | 1.000 | 1.000 |
| CCMpred | 0.203 | 0.276 | 0.490 | 0.583 | 0.150 | 0.203 | 0.286 | 0.333 |
| MetaPSICOV | 0.362 | 0.537 | 0.653 | 0.875 | 0.309 | 0.447 | 0.674 | 0.750 |

**Figure 2.**
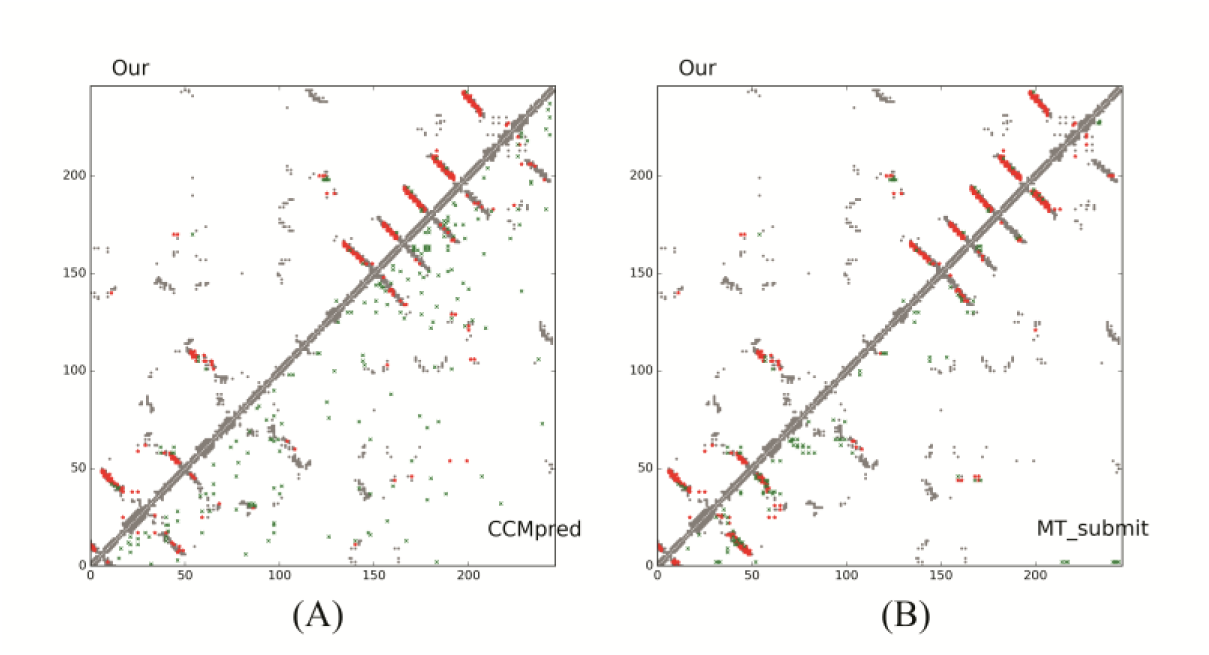
Overlap between predicted contacts (in red and green) and the native (in grey) for T0864-D1. Red (green) dots indicate correct (incorrect) prediction. Top L/2 predicted contacts by each method are shown. (A) The comparison between our prediction (in upper-left triangle) and CCMpred (in lower-right triangle). (B) The comparison between our prediction (in upper-left triangle) and MetaPSICOV-submit (in lower-right triangle).

**Figure 3.**
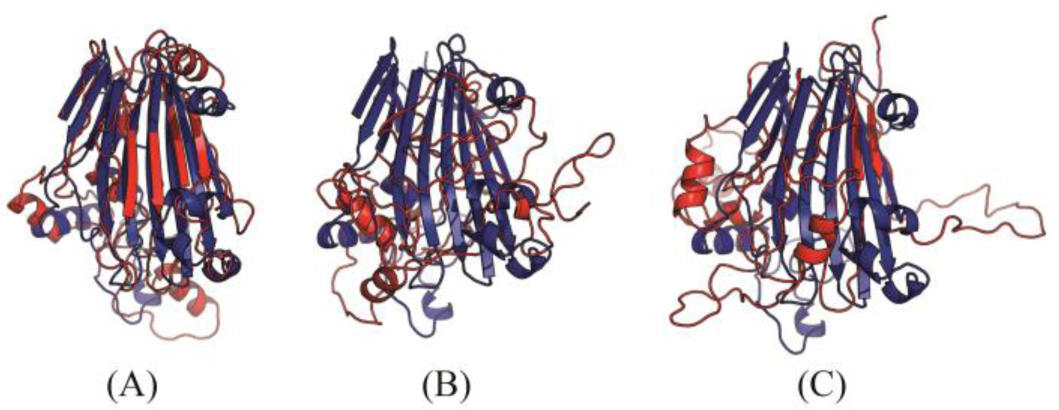
Superimposition between the predicted models (red) and the native structure (blue) for T0864-D1. The models are built by CNS from the contacts predicted by **(A)** our method, **(B)** CCMpred, and **(C)** MetaPSICOV. The TMscores of the three models are 0.63, 0.27 and 0.35, respectively.

### T0869-D1

Table 5 show that our method produced much better contact prediction than CCMpred and MetaPSICOV, especially when top L long-range contacts are evaluated. Specifically, the contact map predicted by our method has top L long-range accuracy 48.1%, while that by CCMpred and MetaPSICOV has corresponding accuracy 10.6% and 23.1%, respectively. Fig. 4 visualizes the top L predicted contacts by the three methods as well as all native contacts. The best of top 5 3D model generated by our method has TMscore 0.69 and RMSD 2.58Å. The best of top 5 3D models built by CNS from CCMpred- and MetaPSICOV-predicted contacts have TMscore 0.265 and 0.441, respectively. Fig. 5 shows the superimposition between the native structure and the 3D models predicted from our method, CCMpred and MetaPSICOV, respectively. To examine the superimposition of our model with its native structure from various angles, please see http://raptorx.uchicago.edu/DeepAlign/25220368/.

**Table 5.** The long- and medium-range contact prediction accuracy of our method, CCMpred, and MetaPSICOV (submit) on the CASP12 target T0869-D1.

|  | Long-range accuracy |  |  |  | Medium-range accuracy |  |  |  |
| --- | --- | --- | --- | --- | --- | --- | --- | --- |
|  | L | L/2 | L/5 | L/10 | L | L/2 | L/5 | L/10 |
| Our method | 0.481 | 0.596 | 0.700 | 0.800 | 0.240 | 0.385 | 0.550 | 0.400 |
| CCMpred | 0.106 | 0.115 | 0.200 | 0.400 | 0.058 | 0.058 | 0.150 | 0.200 |
| MetaPSICOV | 0.231 | 0.308 | 0.500 | 0.600 | 0.125 | 0.192 | 0.200 | 0.300 |

**Figure 4.**
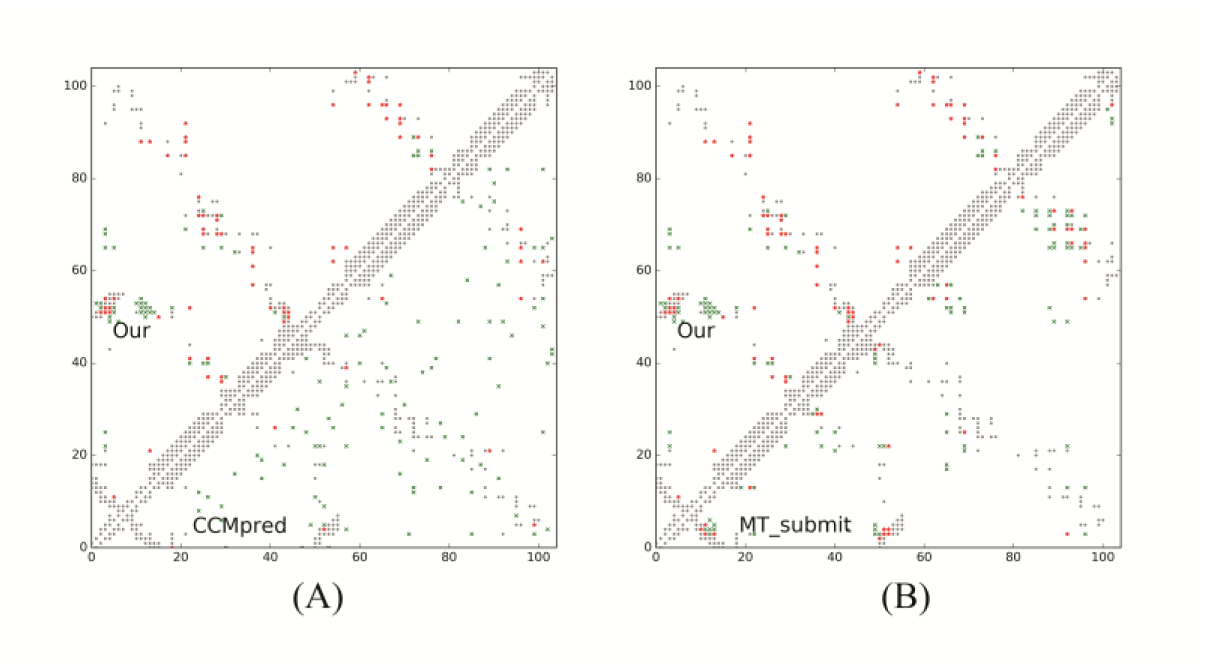
Overlap between predicted contacts (in red and green) and the native (in grey) for T0869-D1. Red (green) dots indicate correct (incorrect) prediction. Top L/2 predicted contacts by each method are shown. (A) The comparison between our prediction (in upper-left triangle) and CCMpred (in lower-right triangle). (B) The comparison between our prediction (in upper-left triangle) and MetaPSICOV (in lower-right triangle).

**Figure 5.**
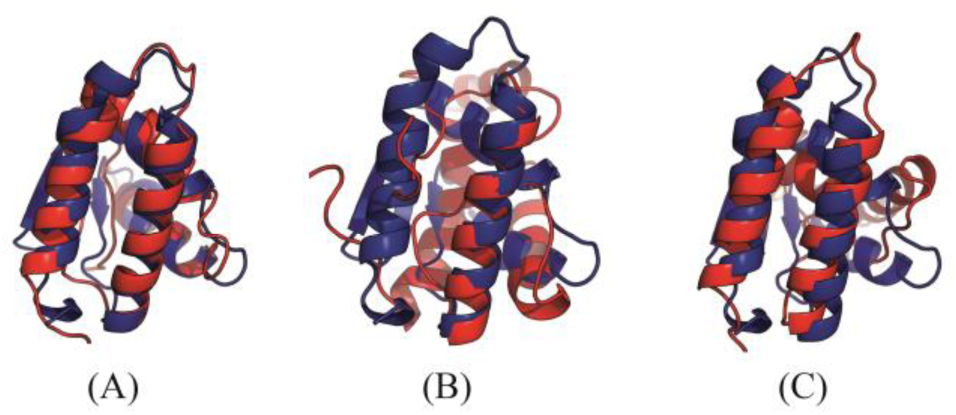
Superimposition between the predicted models (red) and the native structure (blue) for T0869-D1. The models are built by CNS from the contacts predicted by **(A)** our method, **(B)** CCMpred, and **(C)** MetaPSICOV. Their TMscores are 0.690, 0.265 and 0.441, respectively.

### T0904-D1

Table 6 show that our method produced much better contact prediction than CCMpred and MetaPSICOV, especially when top L long-range contacts are evaluated. Specifically, the contact map predicted by our method has top L long-range accuracy 60.6%, while that by CCMpred and MetaPSICOV has corresponding accuracy 4.4% and 17.9%, respectively. Fig. 6 visualizes the top L/2 predicted contacts by each method superimposed to the native contact map. The 3D model generated by our method has TMscore 0.682 and RMSD 4.61Å. The best of top 5 models built by CNS from CCMpred- and MetaPSICOV-predicted contacts have TMscore 0.221 and 0.385, respectively. Fig. 7 shows the superimposition between the native structure and the 3D models generated from our method, CCMpred and MetaPSICOV, respectively. To examine the superimposition of our model with its native structure from various angles, please see http://raptorx.uchicago.edu/DeepAlign/28399687/.

**Table 6.**
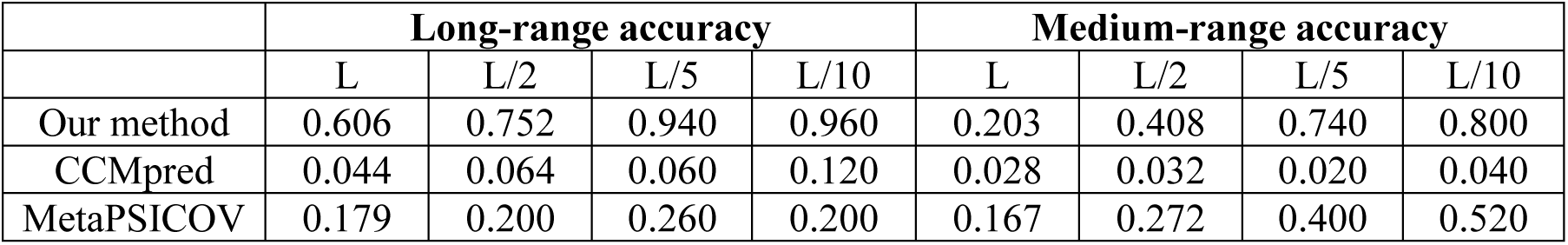
The long- and medium-range contact prediction accuracy of our method, CCMpred, and MetaPSICOV (submit) on the CASP12 target T0904-D1.

|  | Long-range accuracy |  |  |  | Medium-range accuracy |  |  |  |
| --- | --- | --- | --- | --- | --- | --- | --- | --- |
|  | L | L/2 | L/5 | L/10 | L | L/2 | L/5 | L/10 |
| Our method | 0.606 | 0.752 | 0.940 | 0.960 | 0.203 | 0.408 | 0.740 | 0.800 |
| CCMpred | 0.044 | 0.064 | 0.060 | 0.120 | 0.028 | 0.032 | 0.020 | 0.040 |
| MetaPSICOV | 0.179 | 0.200 | 0.260 | 0.200 | 0.167 | 0.272 | 0.400 | 0.520 |

**Figure 6.**
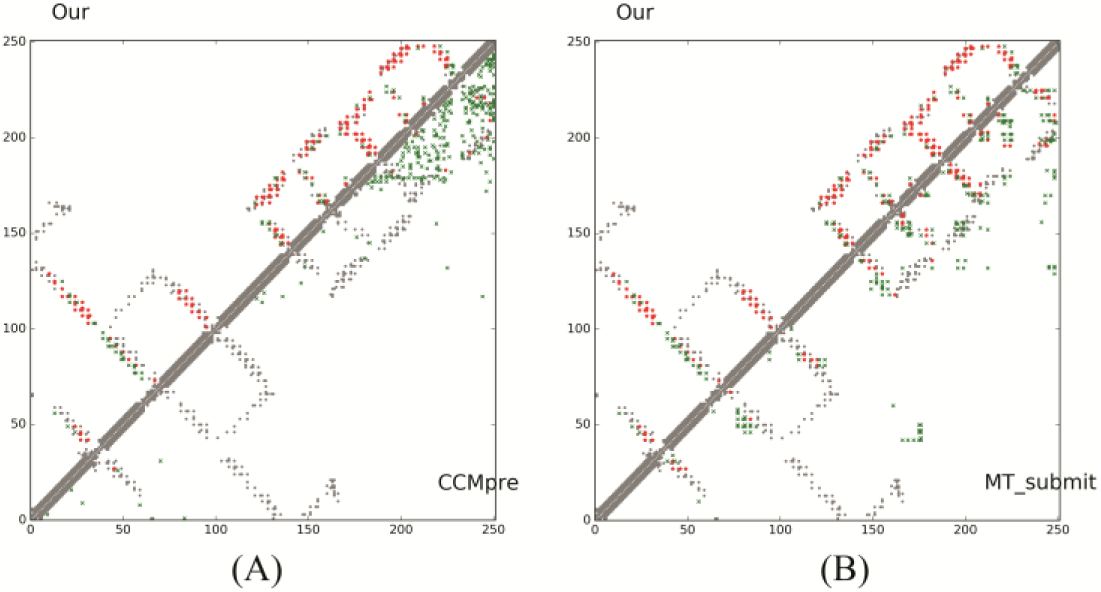
Overlap between predicted contacts (in red and green) and the native (in grey). Red (green) dots indicate correct (incorrect) prediction. Top L/2 predicted contacts by each method are shown. (A) The comparison between our prediction (in upper-left triangle) and CCMpred (in lower-right triangle). (B) The comparison between our prediction (in upper-left triangle) and MetaPSICOV (in lower-right triangle).

**Figure 7.**
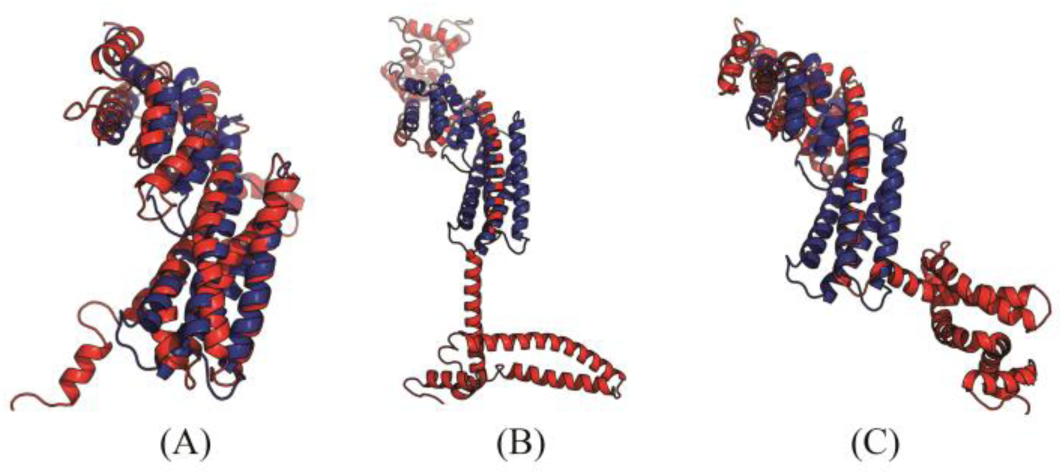
Superimposition between the predicted models (red) and the native structure (blue) for T0904-D1. The models are built by CNS from the contacts predicted by **(A)** our method, **(B)** CCMpred, and **(C)** MetaPSICOV. Their TMscores are 0.682, 0.221 and 0.385, respectively.

## What went right?

Both our self-test results and the CASP12 result indicate that deep learning is a good technique for protein contact prediction although previous deep learning methods such as DNCON [10] and CMAPpro [20] did not stand out in CASP11. This is due to a totally new network architecture we invented. In our in-house benchmark, we found out that increasing the depth of our 2D residual neural network from 1-2 layers to 30 layers can greatly improve contact prediction accuracy. This improvement arises as more layers are used, our learning model can learn more accurately contact occurrence patterns and make the predicted contact maps more protein-like. However, increasing the depth from 30 to 60 layers can only yield a small improvement.

It is possible to obtain good accuracy by using a shallow, but very wide convolutional neural network (i.e., each layer has many more hidden neurons than our current deep model), but we have not rigorously tested such a network yet. One concern with a wide neural network is that to achieve the same performance as our narrow and deep neural network, we may have to use many more hidden neurons at each layer and thus, many more model parameters than our current deep model.

It is important to formulate contact prediction as a pixel-level labeling problem so that we can do simultaneous prediction of all residue pairs in a protein, which allows us to easily capture contact occurrence patterns. For comparison, we developed a deep residual network for the image-level classification formulation of contact prediction. That is, for each residue pair to be predicted, we extract the submatrix (of dimension 41×41) centered around this residue pair, treat it as an image, and assign a positive label to this image if this residue pair forms a contact, otherwise a negative label. Our in-house tests indicate that in terms of top L/10 long-range contact accuracy, image-level classification formulation is about 0.08-0.10 worse than pixel-level labeling formulation. Both formulations use a similar number of 2D convolutional layers and at each layer a similar number of hidden neurons.

## What went wrong?

The major issue is that our method for contact prediction and contact-assisted folding was still under development during CASP12 and thus, we missed a good opportunity to blindly test a fully-implemented deep learning method. Our protocol of building 3D models from predicted contacts was not optimized either. Initially, we fed only top L predicted contacts and predicted secondary structure to CNS[19] to build 3D models. Only at the end of CASP12 did we found out that we should have used more than top L predicted contacts to build better 3D models since our contact prediction had significantly improved.

There are also some other issues in our protocol. For example, we did not handle multi-domain targets very well. For more than half of the multi-domain CASP12 targets, we did not split them into domains for contact prediction. Without domain splitting, sometimes we may miss a good percentage of sequence homologs for some domains (especially those hard targets) and thus, could not generate a good contact prediction. Second, we only employed HHblits and the uniprot20 sequence database dated in February 2016 to generate MSAs while many other predictors also used Jackhmmer[21] and other sequence databases (e.g., metagenomics data), which may be helpful to some targets. Finally, we employed only one tool CCMpred to generate direct co-evolution information while other methods such as MetaPSICOV employed three complementary tools.

## Discussion

We have presented a deep learning method for contact prediction that has much better contact prediction accuracy than existing methods. Our method employs a combination of two deep residual neural networks to model contact occurrence patterns and sequence-contact relationship. As opposed to existing supervised methods that predict contacts of a protein individually, our method predicts all contacts of a protein simultaneously, which makes it easy to model contact occurrence patterns and thus, make predicted contact maps more protein-like. The blind test in CASP12 shows that our deep learning method indeed can do better contact prediction than existing methods. After CASP12, we have been testing our method in a fully-automated, weekly online benchmark CAMEO[22]. The blind test in CAMEO indicates that ab initio folding using our predicted contacts as restraints can fold large proteins without similar structures in PDB and many sequence homologs[13]. It is worth pointing out that currently we fold proteins by feeding predicted contacts to CNS using a very simple protocol. We do not use any fragment assembly, sophisticated energy functions or time-consuming folding simulations. Our folding protocol runs very fast, from 30 minutes to a few hours to generate 200 models on a Linux workstation of 20 CPUs. It is possible to develop a much better folding protocol by combining our predicted contacts with fragment assembly and a sophisticated energy function.

In summary, contact prediction and *ab initio* folding is becoming easier with the advent of direct evolutionary coupling analysis and deep learning techniques. Deep learning can improve contact prediction accuracy not only for proteins with a small number of sequence homologs, but also for proteins with thousands of sequence homologs (see Table 2). We have been studying a few other deep network architectures for protein contact prediction including the dense deep network[23], the wide residual network[24] and the LSTM[25], which have similar or slightly better performance than our current implementation. It is also possible to further improve contact prediction accuracy by including more protein-like features into the deep learning model. For example, the contacts connecting two beta strands of a beta sheet form a special pattern in a contact matrix. If we can incorporate this kind of prior-knowledge into our deep learning model, we shall be able to improve contact prediction accuracy for beta proteins. Nevertheless, predicting contacts for proteins with very few sequence homologs is still very challenging. It is unclear if there is an effective way to use deep learning to yield very accurate contact prediction for this type of proteins.

Our deep learning method also applies to membrane proteins, even though it was not trained on them. We are not aware of any CASP12 hard targets being membrane proteins, but our deep learning method produced decent models for 4 membrane proteins in the blind CAMEO test[26]. This finding implies that the sequence-structure relationship learned by our model from soluble proteins can be transferred to membrane protein contact prediction. Finally, our deep learning method in principle shall also apply to interfacial contact prediction for protein complexes, but may be less effective since on average protein complexes have fewer sequence homologs.

## ACKNOWLEDGMENT

This work is supported by National Institutes of Health grant R01GM089753 to JX and National Science Foundation grant DBI-1564955 to JX. The authors are also grateful to the support of Nvidia Inc. and the computational resources provided by XSEDE through the grant MCB150134 to JX. The funders had no role in study design, data collection and analysis, decision to publish, or preparation of the manuscript. The authors thank CASP12 organizers and assessors and all contributors of the experimental data. We are grateful to Prof. Tobin Sosnick for proofreading our paper.

## Author Contributions

J.X. conceived, designed and implemented the major contact prediction algorithm. S.W. built 3D models from predicted contacts, deployed the algorithm to web server, conducted data preparation and result analysis. Some of S.W.’s data analysis was done while he is affiliated with KAUST, Saudi Arabia. S.S. implemented the deep model for image-level classification formulation. All three authors participated in CASP12 and wrote the paper together.

